# Sequence-aware Prediction of Point Mutation-induced Effects on Protein-Protein Binding Affinity using Deep Learning

**DOI:** 10.1101/2025.11.15.688659

**Authors:** Jianan Zhuang, Zonghui Li, Suhui Wang, Renchao Zheng, Guijun Zhang

**Author notes:** To whom correspondence should be addressed: Guijun Zhang.

## Abstract

Amino acid mutations may lead to significant changes in the binding affinity of protein complexes, thereby causing a series of cellular dysfunctions. Therefore, accurate prediction of protein-protein binding affinity changes (ΔΔG) induced by amino acid mutations is of great importance for understanding protein-protein interactions (PPIs). In this study, we propose SAMAffinity, a protein sequence-aware deep learning architecture for predicting changes in protein-protein binding affinity caused by amino acid mutations. SAMAffinity predicts mutation-induced ΔΔG by integrating multi-source sequence features, leveraging a Mutation-Site Identification (MSI) module to highlight local semantic shifts and a Binding-Interface Awareness (BIA) module to capture interaction changes. Benchmark evaluations on public datasets show that under the mutation-level data splitting strategy, SAMAffinity outperforms the state-of-the-art sequence-based method AttABseq by 33.3%, 72.3%, 31.8%, and 30.5% on S1131, S4169, S645, and M1101 datasets, respectively. Moreover, under the complex-level data splitting strategy, SAMAffinity surpasses the structure-based method MpbPPI by 22.9%, 22.7%, 5.0%, and 11.4% on the corresponding datasets. Beyond predictive accuracy, the strong consistency between the model’s predicted distribution and natural amino-acid mutation tendencies indicates that SAMAffinity effectively captures the underlying mutational landscape shaped by intrinsic biochemical and evolutionary factors. Based on this capability, SAMAffinity demonstrated strong generalization in a study of severe acute respiratory syndrome coronavirus 2 (SARS-CoV-2) cases, suggesting its potential for optimizing therapeutic antibody design.

## INTRODUCTION

Mutation-induced changes in protein-protein binding affinity (ΔΔG) are pivotal for modulating complex interactions and function^1-3^. Even a single amino acid mutation can dictate binding outcomes, directly impacting molecular recognition, signal transduction, and immune responses^4-6^. This is exemplified by antibody affinity maturation, where B cells first diversify through somatic hypermutation^7, 8^, and clones with superior affinity are then selectively expanded to enhance immune efficacy^9^. Consequently, the accurate prediction of ΔΔG is critical for elucidating disease mechanisms and guiding drug and antibody design.

Traditionally, investigations of protein mutational effects relied on wet-lab techniques, notably surface plasmon resonance (SPR)^10^ and isothermal titration calorimetry (ITC)^11^. While these biophysical assays provide precise quantitative measurements, they are often costly and low-throughput^12^. To overcome these limitations, computational methods have been developed. Early representative methods, such as FoldX^13^, Rosetta^14^, and Flex ddG^15^ estimate ΔΔG by modeling protein interactions using physical energy terms. Although theoretically interpretable, these methods are computationally intensive and limited in conformational coverage, which may lead to inaccurate predictions of mutational effects^16^.

To address these limitations, a variety of data-driven methods that leverage structural information have been developed, which can be broadly categorized into two categories. The first emphasizes physicochemical energy terms or structural features combined with traditional machine learning, such as Mutabind2^17^ and TopNetTree^18^. The second focuses on deep learning with graph representations of protein structures, including GeoPPI^19^, MpbPPI^20^, DDMut-PPI^21^ and GearBind^22^, which leverage GNNs^23^ or Transformers^24^ to capture local and hierarchical structural perturbations. Structure-based approaches perform well when high-resolution experimental structural data are available. However, their scalability is limited in large-scale mutation analyses, as each variant requires structure prediction, resulting in substantial computational demands. More critically, protein-protein interactions are inherently dynamic, yet existing models rely on static assumptions, limiting their ability to capture conformational changes upon binding or mutation^25^.

Limitations of structural methods have driven a shift toward sequence-based approaches, which bypass structural modeling by substituting amino acids and comparing embedding changes, enabling high-throughput ΔΔG prediction. In particular, the rapid progress of protein language models^26-29^ has made it possible to extract semantic features such as evolutionary conservation and chemical context from sequences^30^, and several methods now rely solely on sequence data. AttAbseq^31^, for instance, uses attention mechanisms to model antibody-antigen interactions, while DeepInterAware^32^ integrates language model embeddings into a bilinear attention network, both achieving competitive performance without structural inputs. Compared with structure-based methods, sequence-based approaches are faster and more lightweight, but they lack the ability to model three-dimensional conformational changes at binding interfaces and therefore remain less accurate. The fundamental challenge, therefore, lies in enhancing predictive accuracy while maintaining high-throughput efficiency.

In this work, we propose SAMAffinity, a deep neural network model that adopts an encoder-decoder architecture to predict mutation-induced binding affinity changes directly from protein sequences. SAMAffinity leverages Adaptive Feature Convolution (AFC) module as encoder to integrate sequence embeddings from protein language models along with complementary features such as physicochemical properties and evolutionary profiles. The decoder architecture is built around cross-attention mechanisms, comprising two core modules: one is the Mutation-Site Identification (MSI) module and the other is Binding-Interface Awareness (BIA) module. The MSI module specializes in capturing discriminative features between wild-type and mutant-type sequences, while the BIA module is dedicated to highlighting critical residue pairs at the receptor-ligand binding interface. Compared with existing methods, SAMAffinity balances efficiency and robustness, eliminating the need for pre-generated structures and enabling rapid large-scale mutational analysis. In a further evaluation involving SARS-CoV-2 Spike protein and neutralizing antibodies^33, 34^, SAMAffinity’s predictions recapitulated known mutational hotspots and further uncovered a collaborative relationship between specific mutation sites and the associated ΔΔG.

## RESULTS

### An overview of SAMAffinity

SAMAffinity predicts mutation-induced binding affinity changes (ΔΔG) by leveraging an encoder-decoder architecture to capture the contextual dependencies of mutated residues within their sequence environments. As shown in **Figure 1B**, three categories of features are extracted from the input sequences: (i) physicochemical properties, such as polarizability, secondary structure^35^, and relative accessible surface area (RASA)^36^, which characterize the local microchemical environment of amino acids; (ii) evolutionary profiles, including the BLOSUM62^37^ substitution matrix and the position-specific scoring matrix (PSSM)^38^, reflecting the evolutionary conservation of amino acid sequences; and (iii) protein language model embeddings, derived from ESM-2^28^ and ProtT5^27^, to capture the residue-level contextual semantic information. In particular, for antibody sequences, the ProtT5 embeddings are replaced with AntiBERTy^26^ embeddings, for which was specifically trained on antibody datasets that better captures the affinity maturation trajectory of antibodies. Details and descriptions of all features are provided in the **Methods** section, as well as in **Supplementary Tables S1 and S2**.

**Fig. 1.**
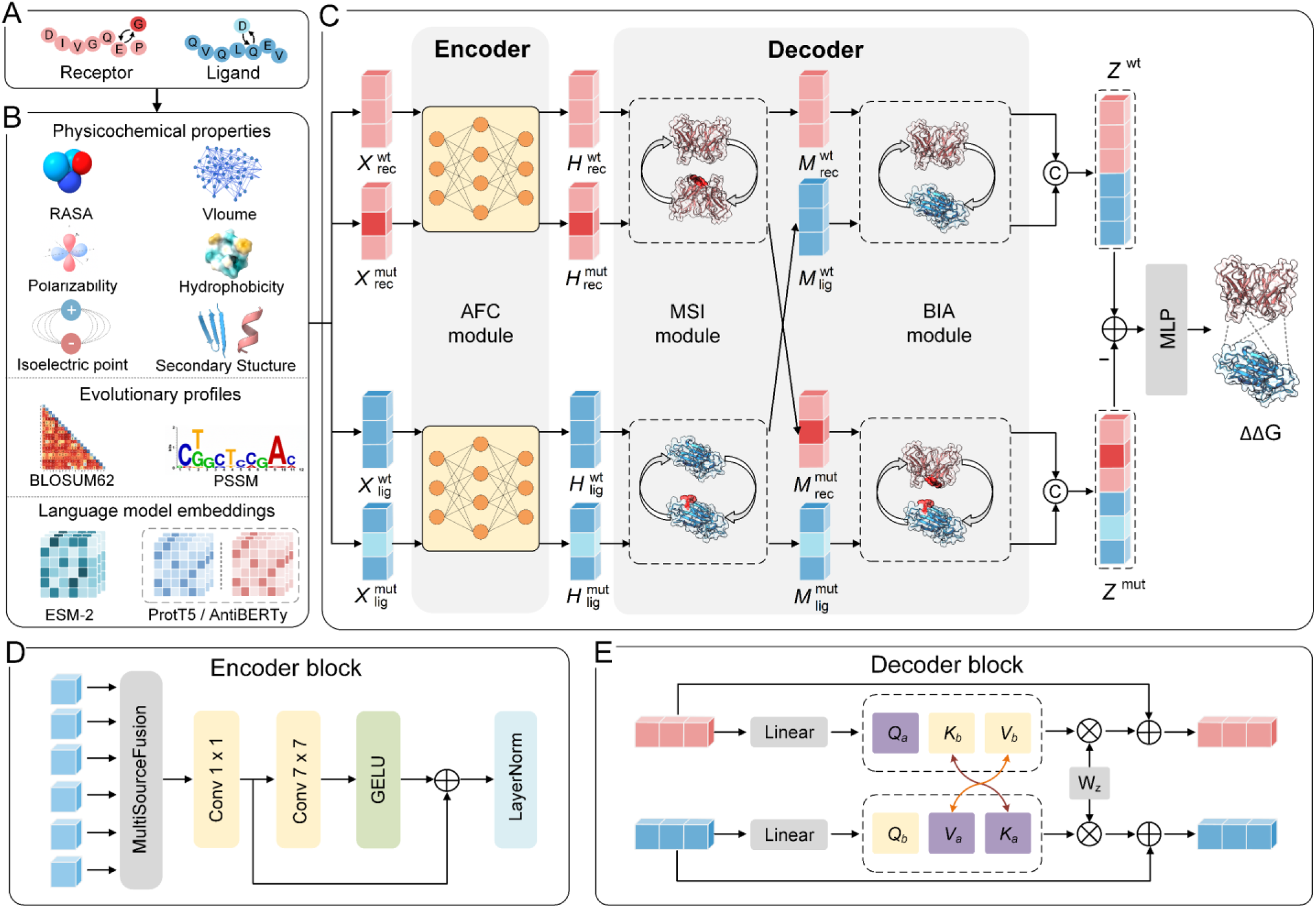
Pipeline of SAMAffinity. (A) The input protein complex sequence is split into receptor and ligand sequences, and corresponding mutant sequences are generated according to the mutation information. (B) Three categories of features are extracted from the input sequences: physicochemical properties, evolutionary profiles, and language model embeddings. Specifically, the physicochemical properties comprise relative accessible surface area (RASA), volume, polarizability, hydrophobicity, isoelectric point, and secondary structure. The evolutionary profiles include BLOSUM62 and the position-specific scoring matrix (PSSM), while the language model embeddings are derived from ESM2 and ProtT5. For antibody-antigen complexes, the ProtT5 embedding of the antibody is replaced by AntiBERTy. (C) Network Architecture: input features are divided into four components: receptor-wild-type, receptor-mutant-type, ligand-wild-type and ligand-mutant-type. The model takes these four components as input, employing an Adaptive Feature Convolution (AFC) module to integrate multiple features. It further adopts a decoder composed of the Mutation-Site Identification (MSI) module and Binding-Interface Awareness (BIA) module to learn mutation effects and protein-protein interactions, respectively. Finally, the model performs differential concatenation between the obtained wild-type and mutant-type feature vectors, and then passes the result through a fully connected layer to reduce dimensionality and output the predicted ΔΔG value. (D) Design of encoder block, MultiSourceFusion layer represents independent linear layers which assign weights to each feature. (E) Design of decoder block, this block accepts a pair of inputs, either wild/mutant or receptor/ligand, and outputs the corresponding updated vectors.

The overall network architecture is shown in **Figure 1C**. All input feature tensors are divided into four components: receptor-wild-type, receptor-mutant-type, ligand-wild-type, and ligand-mutant-type. The receptor and ligand feature tensors are independently processed through two pairs of encoder-decoder networks with non-shared parameters. As illustrated in **Figure 1D**, the encoder is composed of AFC module, in which each feature is assigned an independent linear layer for adaptive weighting. The weighted features are subsequently concatenated, and a 1D convolutional block is applied to extract high-dimensional local representations. In the decoder **(Fig. 1E)**, the wild-type and mutant-type representations of the same chain first interact within the MSI module, allowing the model to capture tensor perturbations caused by amino acid substitutions. Thereafter, the BIA module enables information exchange between receptor and ligand representations, thereby facilitating the learning of interaction patterns underlying receptor-ligand binding. Detailed algorithms are provided in the **Methods** section.

In the downstream output stage, residue-level features are aggregated into sequence-level representations using a combination of Softmax normalization and summation pooling. The ΔΔG computation is explicitly modeled by performing an element-wise subtraction of the mutant-type tensor from the wild-type tensor. This difference tensor is then passed through a fully connected layer to output the final predicted ΔΔG value.

### Mutation-level correlation comparison

To fairly evaluate the performance of SAMAffinity, we selected four widely recognized public benchmark datasets: S1131^39^, S4169^40^, S645^18^ and M1101^41^. Among them, S1131 and S4169 are single-point mutation datasets derived from SKEMPI 2.0^42^, S645 is a single-point mutation dataset from AB-Bind^41^, and M1101 represents the full AB-Bind dataset comprising both single and multiple point mutations. For experimental design, we employed a 10-fold cross-validation strategy, where mutations were randomly shuffled and partitioned to ensure balanced distribution. The linear correlation of the model’s outputs was primarily evaluated using the Pearson correlation coefficient. In addition, MAE and RMSE were employed to assess the prediction error.

To establish a comprehensive benchmark, SAMAffinity was evaluated against both the state-of-the-art sequence-based approaches AttABSeq^31^ and structure-based methods MpbPPI^20^. This dual comparison is essential because structure-based methods are often considered gold standards, leveraging physicochemical and spatial information, whereas sequence-based approaches like SAMAffinity aim to achieve competitive performance using only primary sequence data. Thus, the contrast rigorously tests whether SAMAffinity can effectively bridge the information gap between sequence and structure.

Since AttABSeq did not report results on the S4169 dataset, we reproduced its outcomes under our experimental conditions to ensure a fair comparison. Results for all other baselines were taken directly from their original publications.

On the two SKEMPI 2.0^42^ subsets, S1131 **(Fig. 2A)** and S4169 **(Fig. 2B)**, SAMAffinity demonstrated robust predictive performance, with data points closely distributed along the diagonal line, indicating strong agreement between predicted and experimental values. Specifically, the Pearson correlation coefficients reached 0.88 **(Fig. 2E)** and 0.81 **(Fig. 2F)** on the two datasets, respectively, which not only significantly outperformed the state-of-the-art sequence-based method AttABseq, with improvements of 33.3% (paired Steiger’s Z-test, Z=24.21, *P* < 0.001) and 72.3% (paired Steiger’s Z-test, Z=51.51, *P* < 0.001), but also slightly surpassed the best structure-based method MpbPPI. In addition, compared with AttABseq, SAMAffinity exhibited consistently lower prediction errors across datasets. On the S1131 dataset, the MAE and RMSE were reduced by 31.7% (paired Wilcoxon signed-rank test, *P* = 4.92×10^-47^) and 35.7% (paired Wilcoxon signed-rank test, *P* = 1.51×10^-45^), respectively. Similarly, on the S4169 dataset, the MAE and RMSE were reduced by 34.0% (paired Wilcoxon signed-rank test, *P* = 5.05×10^-273^) and 31.6% (paired Wilcoxon signed-rank test, *P* = 7.34×10^-257^), respectively. These consistent improvements across different datasets demonstrate that SAMAffinity yields smaller prediction deviations and higher numerical precision in estimating binding affinities. The extremely narrow 95% confidence intervals further confirmed the stability and robustness of the model’s predictions under large-sample conditions.

**Fig. 2.**
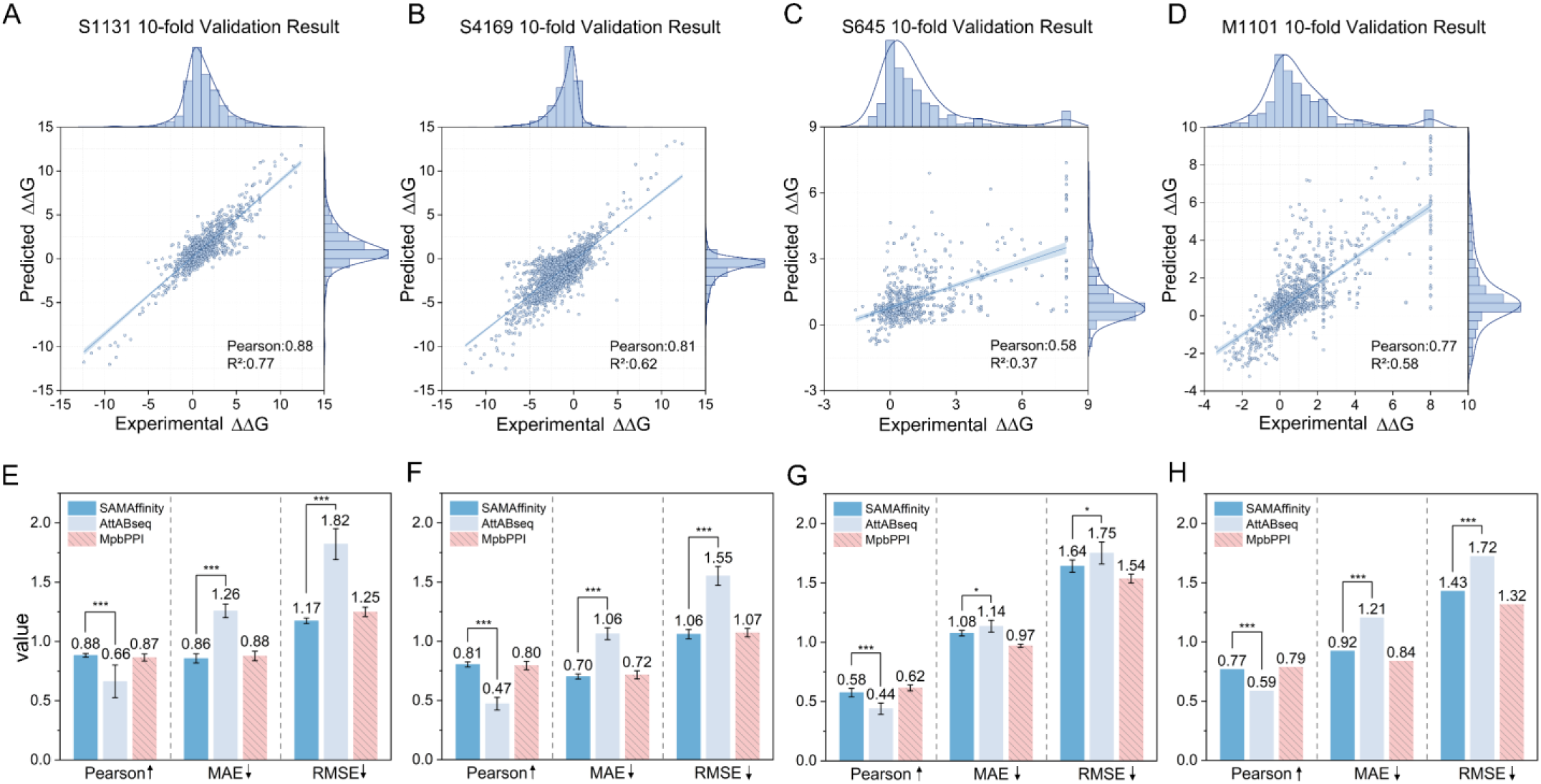
Comparison between SAMAffinity with other methods on the four benchmark datasets under mutation-level 10-fold cross-validation. (A-D) show the scatter plots of SAMAffinity on each dataset, the frequency distribution histograms on the top and right sides of the scatter plot respectively illustrate the data distributions of the true labels and the predicted results. Light colored shadows indicate a 95% confidence interval, estimated using the Student’s t-test. (E-H) show the mean comparison results for Pearson, MAE, and RMSE, with error bars representing the standard deviation (SD). The hatched pattern indicates that the method is structure-based, and the direction of the arrows next to each metric denotes the direction of improvement. The statistical significance of differences in Pearson correlation coefficients between SAMAffinity and AttABseq predictions was assessed using Steiger’s Z-test for paired correlations within the same sample set. For error metrics (MAE and RMSE), the normality of paired differences was first evaluated using the Shapiro-Wilk test. Two-sided paired t-tests were applied to normally distributed differences, while non-normally distributed differences were evaluated using two-sided Wilcoxon signed-rank tests. Significance levels are denoted as P ≥ 0.05 (not significant), P < 0.05 (*), P < 0.01 (**), and P < 0.001 (***). Due to the unavailability of sample-level data for MpbPPI, significance analyses involving this method were not performed.

On the two AB-Bind subsets, S645 **(Fig. 2G)** and M1101 **(Fig. 2H)**, SAMAffinity achieved Pearson correlation coefficients of 0.58 and 0.77, showing relative increases of 31.8% (paired Steiger’s Z-test, Z=5.13, *P* < 0.001) and 30.5% (paired Steiger’s Z-test, Z=7.15, *P* < 0.001) over AttABseq, respectively. For the MAE and RMSE metrics, though the reductions are not as pronounced as those observed in the two subsets of SKEMPI 2.0, they still decreased by 5.3% (paired Wilcoxon signed-rank test, *P* = 1.88×10^-2^) and 6.3% (paired Wilcoxon signed-rank test, *P* = 1.23×10^-2^) on the S645 dataset. On the M1101 dataset, the decreases were 24.0% (paired Wilcoxon signed-rank test, *P* = 1.63×10^-10^) and 16.9% (paired Wilcoxon signed-rank test, *P* = 7.55×10^-10^), respectively. Although on the two AB-Bind subsets, SAMAffinity did not surpass the best structure-based method MpbPPI, it still maintained competitive performance. It should be noted that the scatter plots **(Fig. 2C and D)** exhibited a significant number of outliers, wherein a cluster of data points corresponded to a fixed experimental value of 8. These outliers are caused by inherent noise in the AB-Bind: when a mutation leads to complex dissociation, the corresponding dissociation constant exceeds the measurable range, and AB-Bind assigns a fixed value of 8 to these samples. Since these truncated values fail to reflect the true strength of mutation effects, they introduce noise into model fitting and reduce correlation in predictions. Details of the significance analysis methods are provided in the **Supplementary Note S1.**

Overall, SAMAffinity exhibited distinct performance patterns across the two datasets. On the SKEMPI 2.0 subsets, SAMAffinity achieved larger improvements compared with AttABseq and slightly outperformed the structure-based method MpbPPI. In contrast, on the AB-Bind subsets, the performance gains were more modest, and the results were slightly worse than MpbPPI. We speculate that this performance gap primarily stems from the intrinsic characteristics of the datasets. SKEMPI 2.0 contains diverse protein-protein complexes with a wide range of interface types and mutation effects, providing a broader spectrum of binding affinity variations. By comparison, AB-Bind focuses exclusively on antibody-antigen complexes, which tend to present more specialized and highly constrained binding interfaces. These dataset-specific factors account for the observed variation in performance, while SAMAffinity consistently demonstrates stable and robust predictive capabilities across both general and antibody-antigen protein complexes.

### Complex-level correlation comparison

Numerous studies have shown that mutation-level cross-validation can lead to overly optimistic performance estimates due to data leakage between training and test sets. To mitigate this bias and rigorously assess generalizability, we implemented a complex-level 5-fold cross-validation scheme, consistent with the strategy employed in MpbPPI^20^ and related works. In short, during data splitting, we ensured that mutation samples derived from the same wild-type complex were exclusively assigned to either the training or validation set, thereby preventing evaluation bias caused by data similarity. Detailed design of our data splitting strategy is provided in the **Supplementary Note S2**.

Under the complex-level data splitting strategy, SAMAffinity achieved the highest Pearson correlation coefficients across all four datasets. Specifically, the model attained correlation values of 0.59 (**Fig. 3A**, S1131), 0.54 (**Fig. 3B**, S4169), 0.42 (**Fig. 3C**, S645), and 0.49 (**Fig. 3D**, M1101), which represent relative improvements of 51.3% (paired Steiger’s Z-test, Z=14.35, *P* < 0.001), 45.9% (paired Steiger’s Z-test, Z=32.66, *P* < 0.001), 55.6% (paired Steiger’s Z-test, Z=5.04, *P* < 0.001) and 63.3% (paired Steiger’s Z-test, Z=7.82, *P* < 0.001) over the sequence-based method AttABseq, respectively. Moreover, SAMAffinity also outperformed the structure-based method MpbPPI, with improvements of 22.9%, 22.7%, 5.0% and 11.4% on the corresponding datasets. For the regression error metrics, SAMAffinity achieved lower MAE and RMSE values than MpbPPI on both SKEMPI 2.0 subsets, with reductions of 13.0% and 13.5% on S1131, and 4.7% and 4.7% on S4169, respectively. However, on the two AB-Bind subsets, S645 and M1101, its error values were slightly higher than those of MpbPPI.

**Fig. 3.**
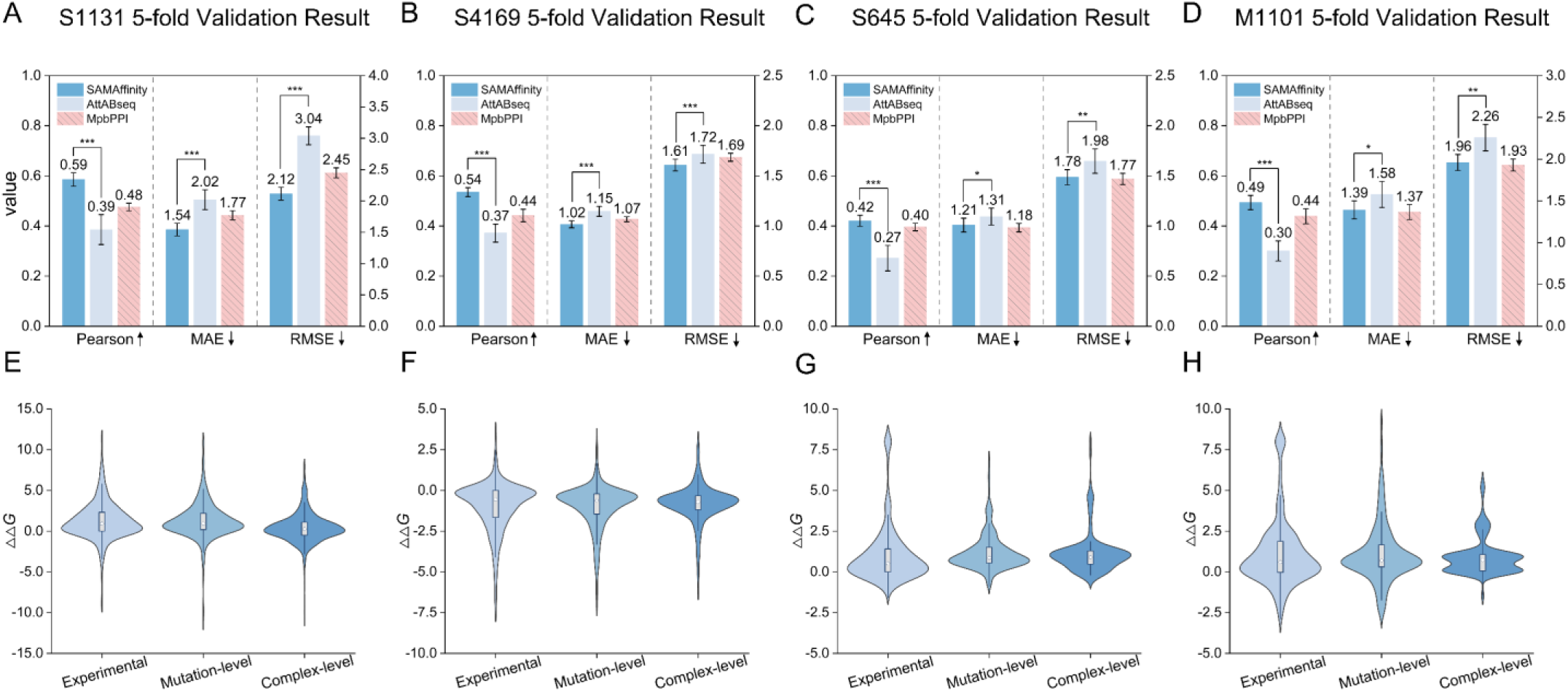
(A-D) Comparison of SAMAffinity with other methods on four benchmark datasets under complex-level 5-fold cross-validation. Pearson correlation coefficients correspond to the left y-axis, while MAE and RMSE values correspond to the right y-axis. The hatched pattern indicates that the method is structure-based, and the direction of the arrows next to each metric denotes the direction of improvement. Error bars represent standard deviation (SD). The significance level test was conducted under the same settings as the mutation-level split strategy. (E-H) Violin plots show the predictive stability of SAMAffinity under different data splitting strategies, the box represents quartiles, the whiskers indicate outliers.

The divergence between correlation-based and error-based metrics reflects intrinsic differences in model generalization and calibration. Under the mutation-level split strategy, structural information dominates and mitigate performance gaps, as data leakage allows both models to access similar complex-level information. In contrast, the complex-level split enforces strict generalization to unseen complexes, where sequence-based models like SAMAffinity better capture transferable statistical patterns, achieving higher correlations. However, the lack of explicit structural constraints leads to less accurate absolute ΔΔG estimation, yielding higher MAE and RMSE, particularly in datasets with narrow energetic ranges such as AB-Bind. This highlights that SAMAffinity effectively learns the directional trends of binding affinity changes, while structure-based methods retain advantages in scale calibration.

To more finely evaluate the differences in prediction distribution caused by the two data splitting strategies, we compared the distributions of the true and predicted ΔΔG values under the mutation-level and complex-level settings using violin plots. On the S1131 (**Fig. 3E**) and S4169 (**Fig. 3F**) datasets, the model exhibited high consistency across both data splitting strategies. While the predicted distribution under complex-level showed a slight shift relative to mutation-level, the overall distribution shapes remained largely preserved, indicating SAMAffinity’s robustness under different data splitting strategies. In contrast, the predicted distribution of the S645 dataset (**Fig. 3G**) exhibits a more pronounced density peak shift under the complex-level setting, whereas for the M1101 dataset (**Fig. 3H**), it changes directly from a unimodal to a bimodal distribution. This variation may arise from the distinctiveness of antibody-antigen complexes: antibody mutations lead to broader and more heterogeneous changes in binding affinity. Under strict complex-level splitting, although SAMAffinity achieves higher trend consistency, the limited data and the amplification effect of certain extreme samples may lead to prediction biases.

Overall, the cross-validation results at the complex-level indicate that SAMAffinity is better at learning the trend of mutational effects rather than predicting their precise scale. In practical protein design scenarios, the trend of mutational effects is often more important, as it indicates whether an amino acid substitution is positive or negative. Due to the inherent evolutionary conservation in sequence data, SAMAffinity demonstrates more robust and stable generalization ability. In contrast, structure-based ΔΔG prediction methods are limited by the accuracy of structural modeling and often struggle to maintain stable performance on new data. Moreover, protein-protein interactions are inherently dynamic, while most structural modeling approaches focus on static conformations, limiting their ability to capture conformational changes during binding process. Sequence-based methods, by avoiding reliance on structural data, can partially mitigate these limitations.

### Feature ablation

Using the optimal model identified via 10-fold cross-validation, we performed ablation studies on the four benchmark datasets to evaluate the contribution of the selected features. Their effectiveness was quantified using Pearson correlation coefficient. Notably, in the S4169 dataset, some samples contain fewer than 10 residues, making the calculation of RASA and PSSM features infeasible. For these samples, zeros were filled in the corresponding feature channels to ensure consistency in feature dimensions. For the antibody-antigen interaction datasets (M1101 and S645), we additionally assessed the utility of antibody sequence embeddings derived from the AntiBERTy.

Ablation analysis across four datasets highlight the distinct contributions of language model embeddings, evolutionary profiles, and physicochemical properties. As illustrated in **Figure 4A**, S1131 and S4169 show substantial drops in Pearson correlation when ESM2 or ProtT5 embeddings are removed (from 0.883, 0.813 to 0.853/0.843 and 0.762/0.778). These embeddings are designed to capture contextual dependencies within protein sequences, reflecting long-range residue interactions and evolutionary regularities learned from massive sequence corpora. Their pronounced impact underscores that language-model-derived representations provide a rich, generalized encoding of residue environments and mutational sensitivities that cannot be fully recovered by handcrafted descriptors. Similarly, in **Figure 4B**, antibody-antigen datasets S645 and M1101 exhibit marked declines when AntiBERTy embedding is excluded (from 0.590/0.768 to 0.542/0.735), highlighting the model’s specialization for the unique compositional and structural characteristics of antibody variable regions, which is capable of effectively capturing the affinity maturation trajectory of antibodies.

**Fig. 4.**
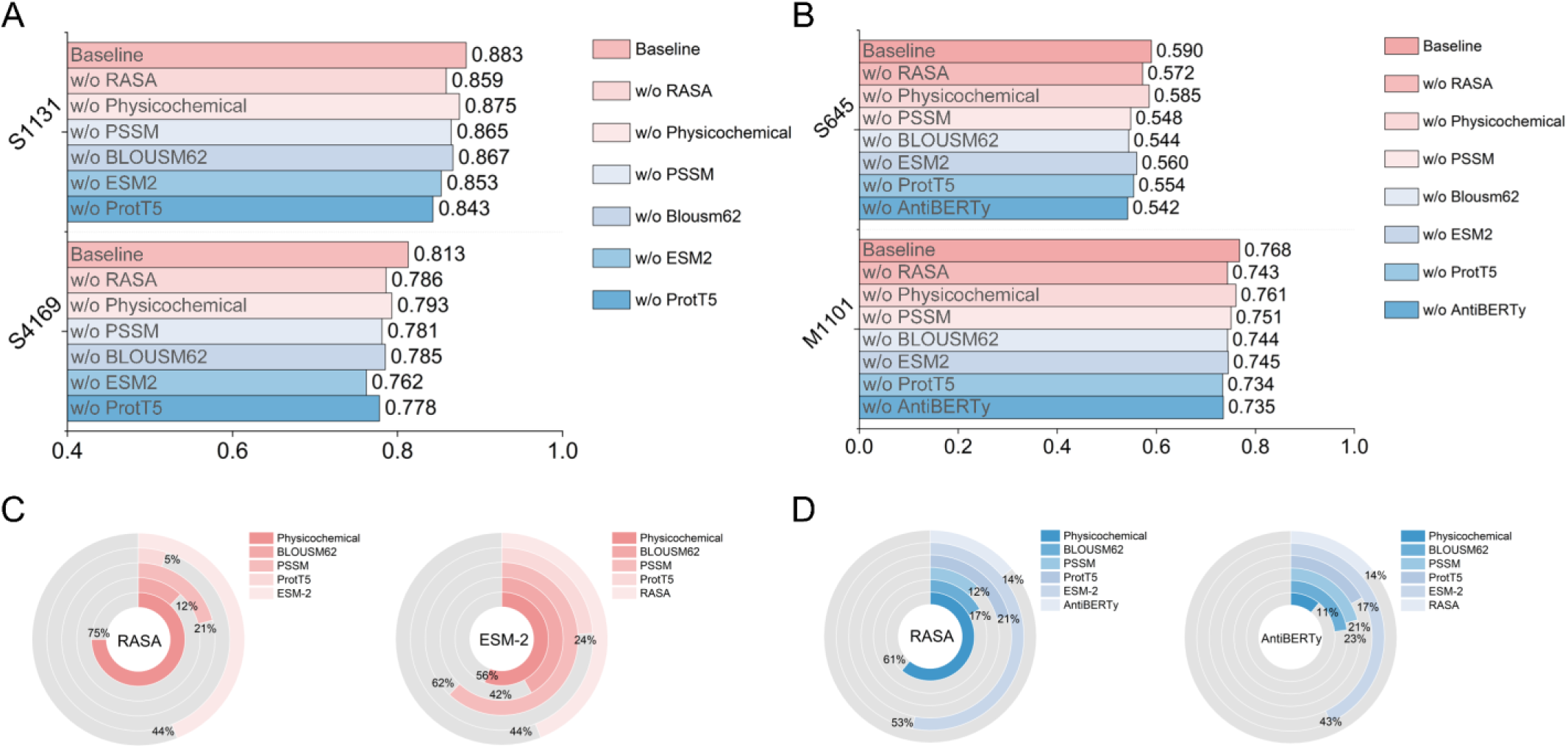
10-fold cross-validation ablation results on four benchmark datasets. “w/o” indicates the exclusion of a specific feature. (A) Ablation results on the two SKEMPI 2.0 subsets. (B) Ablation results on the two AB-Bind subsets, with the addition of the AntiBERTy embedding. (C) Correlations between RASA, ESM-2 and other features measured on the two SKEMPI 2.0 subsets. (D) Correlations between RASA, AntiBERTy and other features measured on the two AB-Bind subsets.

Evolutionary profiles (BLOSUM62 and PSSM) also made pronounced contributions, especially in S645 and M1101. These features were designed to encode residue-level substitution propensities and conservation scores derived from multiple sequence alignments. Their strong impact indicates that evolutionary constraints remain vital for interpreting the biological significance of mutations-providing explicit, human-interpretable signals that complement the implicit evolutionary priors already encoded in large-scale language model embeddings.

Finally, physicochemical properties enhance the model’s interpretability by explicitly encoding charge, hydrophobicity, and other local environmental descriptors, thereby bridging the gap between abstract embeddings and interpretable biochemical attributes. In particular, the relative accessible surface area (RASA) consistently improved performance across all four datasets, reflecting its ability to characterize biophysical exposure and residue accessibility, which are key determinants of interaction energetics and stability upon mutation. By quantifying how buried or exposed a residue is, this feature provides essential physical context that purely sequence-based embeddings may overlook.

To verify the necessity of the selected features, we conducted feature correlation analyses on the SKEMPI 2.0 and AB-Bind datasets, respectively. In the SKEMPI 2.0 dataset **(Fig. 4C)**, although RASA exhibited high redundancy with physicochemical properties, its correlation with other features was relatively low. In contrast, ESM-2 showed moderate redundancy with all other features. Notably, the low redundancy between ESM-2 and ProtT5 highlights their distinct feature-capturing mechanisms: ESM-2 focuses more on extracting evolutionary information from amino acid sequences, whereas ProtT5 emphasizes the functional semantics of residues within their contextual environment. Similarly, in the AB-Bind dataset **(Fig. 4D)**, RASA exhibited a correlation pattern consistent with that observed in SKEMPI 2.0. In contrast, AntiBERTy, as an antibody-specific language model, showed low correlations with all other features, highlighting its distinct representation space and explaining its superior performance observed in the ablation experiments.

The feature correlation analyses further confirm the necessity of the multi-source feature integration strategy: language model embeddings provide global sequence dependency information, evolutionary profiles capture evolutionary constraints, and physicochemical properties characterize the local microchemical environment. These complementary features collectively enhance the model’s robustness and generalization capability.

### Benchmarking predictive models against natural amino acid mutation trends

Amino acid mutations in nature are not entirely random, rather, they are shaped by a combination of physicochemical properties, functional constraints, and evolutionary pressures, resulting in discernible patterns. In this section, we examine the consistency between the model-predicted and experimentally observed distributions of average binding affinity changes across different amino acid substitutions. Specifically, based on the S1131 dataset, we systematically analyzed the consistency between the actual mutational patterns of the 20 amino acids **(Fig. 5A)** and the model’s predicted patterns **(Fig. 5B)**, and quantitatively assessed their alignment by calculating the distribution of differences between the two **(Fig. 5C)**.

**Fig. 5.**
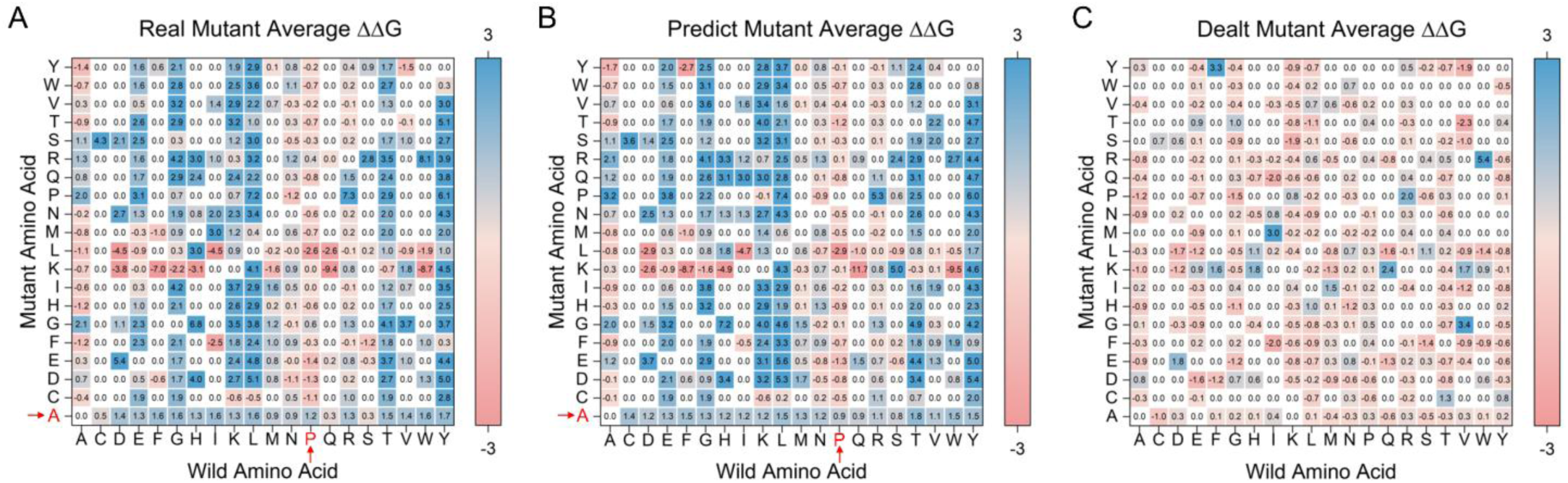
Consistency analysis of mutational tendencies predicted by SAMAffinity on the S1131 dataset. (A) Distribution of average binding affinity changes for amino acid mutations, computed from the true labels in the dataset. The x-axis represents the wild-type amino acid, and the y-axis represents the mutant amino acid. Each point denotes the average ΔΔG value for a specific mutation type; white points (value = 0) represent mutation types with no corresponding samples in the dataset. (B) Distribution of average binding affinity changes computed from SAMAffinity’s predicted results. (C) Difference between the true and predicted distributions, calculated as Δ = true_value - predicted_value.

The heatmap analysis reveals a strong consistency between the model-predicted and experimentally observed distributions of average binding affinity changes. Interestingly, we observed that mutations involving proline (Pro) tend to result in a general increase in binding affinity. This trend may be attributed to the unique structural properties of proline: its α-carbon side chain forms a pyrrolidine ring by covalently bonding to the backbone nitrogen, converting the typical α-amino group (-NH2) into a secondary amine group (-NH-). This cyclic structure imparts greater rigidity to proline and often disrupts ordered secondary structures, thereby enhancing flexibility in disordered regions. However, this special structural feature also limits proline’s ability to form peptide bonds and prevents it from serving as a hydrogen bond donor^43^. As a result, proline is generally less involved in high-affinity interaction interfaces, and its substitution may, in some cases, relieve structural constraints, leading to improved binding affinity.

Another noteworthy observation is that when other amino acids are mutated to alanine (Ala), the binding affinity between proteins tends to decrease consistently. In the field of protein-protein interaction research, alanine scanning mutagenesis^44^ is a well-established strategy. This approach leverages the small, chemically inert methyl side chain of alanine to systematically evaluate the functional importance of interface residues. Typically, substituting a residue with alanine leads to a reduction in binding affinity, as the mutation often disrupts critical side-chain interactions, which is particularly pronounced at hotspot residues.

Furthermore, **Figure 5C** shows that the overall distribution of differences exhibits a slight negative shift, indicating a positive bias in the model’s predictions relative to the experimental distribution. This phenomenon can be attributed to the inherent label distribution of the S1131 dataset, which is skewed toward positive values. As a result, the model tends to internalize this distributional bias during training, leading to a systematic overestimation in its predicted ΔΔG values.

### SARS-CoV-2 case studies

To further evaluate model performance, we compiled a test set of 70 single-point mutations from SARS-CoV-2-antibody binding interfaces, sourced from the literature^33, 34^. We excluded closely related homologs from the training data using a 99% antibody sequence identity threshold and ensured temporal truncation between the test set and the training data. (See **Supplementary Table S3** for the full test set composition) For comparison, we selected the structure-based predictor, DDMut-PPI’s web server, as a baseline. Since DDMut-PPI^21^ is a structure-based predictor, we aimed to simulate a challenging and realistic ‘blind-test’ scenario where experimental structures are unavailable. Therefore, we used the AlphaFold 3^45^ server to predict the wild-type complex structures, which then served as input for DDMut-PPI. This setup rigorously evaluates the performance in practical applications.

As shown in **Figure 6A** and **6B**, SAMAffinity achieves a Pearson correlation of 0.473 on the test set, representing a substantial improvement over DDMut-PPI’s 0.314. This difference not only indicates stronger overall fitting ability of SAMAffinity but also reflects its greater robustness in capturing the linear correlation between experimental ΔΔG values and predicted results. From the fitted curves and the 95% confidence intervals, it is evident that SAMAffinity’s predictions are more concentrated, with most points lying close to the diagonal line, whereas DDMut-PPI exhibits greater dispersion, suggesting lower stability in handling certain mutations.

**Fig. 6.**
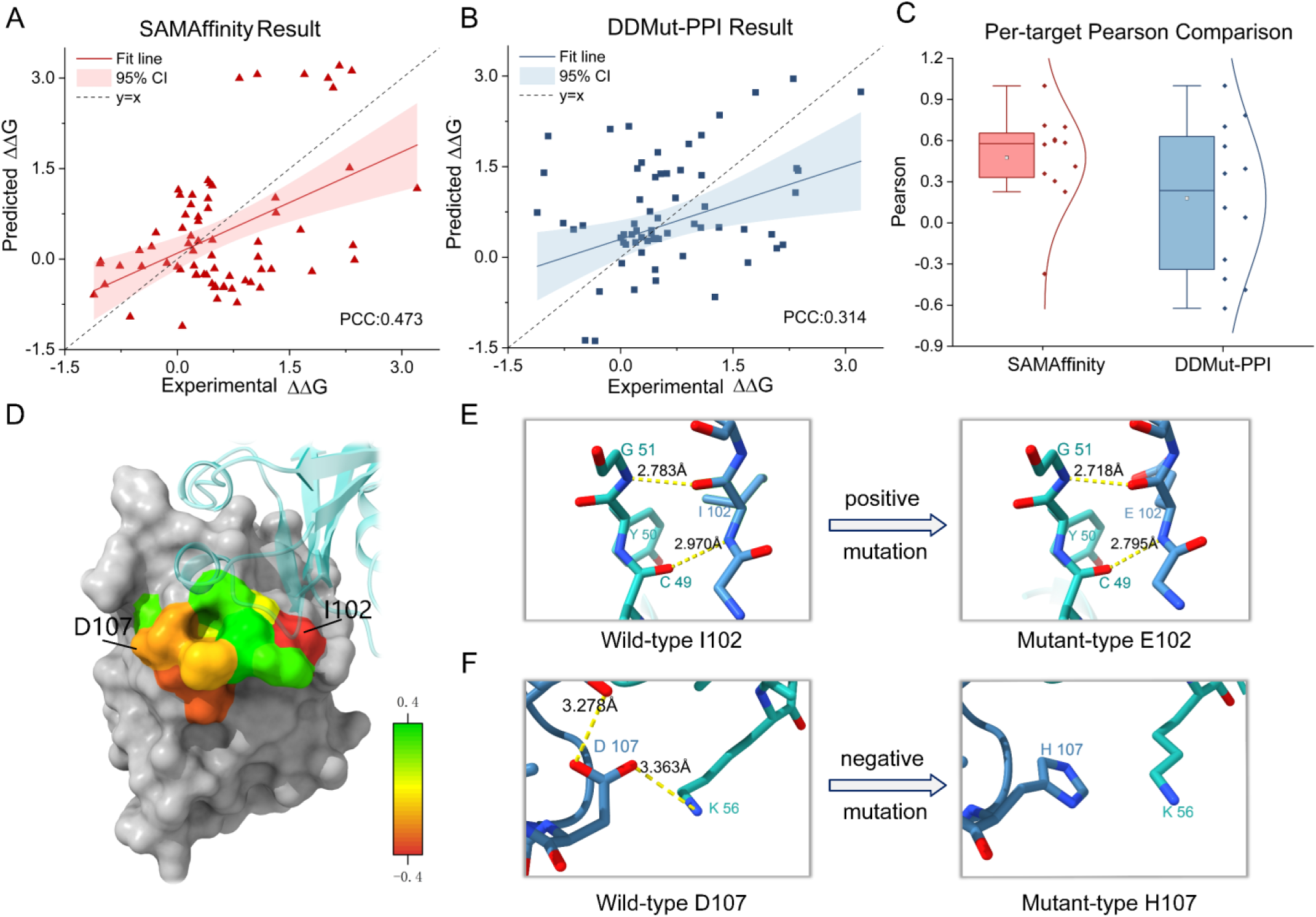
Case study results on SARS-CoV-2. (A) Scatter plot of SAMAffinity predictions on the SARS-CoV-2 test set, with a Pearson correlation coefficient of 0.473. The gray dashed line represents y = x, the red line indicates linear regression, and the shaded red area shows the 95% confidence interval. (B) Scatter plot of DDMut-PPI predictions on the same test set, with a Pearson correlation coefficient of 0.314. (C) Comparison of per-target Pearson correlation coefficients between SAMAffinity and DDMut-PPI. Horizontal lines indicate the median; white boxes represent the mean values. (D) Heatmap of average predicted binding affinity changes at each residue position in the CDR-H3 region of CR3022. Two key mutation sites were identified: H: I102E and H: D107H, represent the most positive and negative mutation site, respectively. (E) AlphaFold 3 structural analysis of the H: I102E mutation. Left: wild-type I102; right: mutant-type E102. (F) AlphaFold 3 structural analysis of the H: D107H mutation. Left: wild-type D107; right: mutant-type H107.

Furthermore, **Figure 6C** illustrates the per-target Pearson distributions across different antibody-antigen complexes. It can be observed that SAMAffinity maintains high consistency across most complexes, with median values in the boxplots significantly higher than those of DDMut-PPI and narrower ranges of variation. These results indicate that SAMAffinity generalizes better across complexes, effectively learning and transferring mutational patterns across different antibody-antigen pairs. It is noteworthy that SAMAffinity’s individual data points are more evenly distributed, with no extreme low values, demonstrating its cross-target consistency and robustness.

In addition, we attempted to use SAMAffinity for CDR-H3 region design of the CR3022 antibody. We adopted a systematic scanning strategy to perform single-point substitutions of all 20 amino acid types at each position within the CDR-H3 region of the CR3022 antibody. The binding affinity changes (ΔΔG) of each mutant relative to the wild-type was predicted using SAMAffinity. By ranking the predicted ΔΔG values and visualizing the results as a heatmap **(Fig. 6D)**, we successfully identified two potential key mutation sites, H: I102E and H: D107H, which exhibited the most pronounced enhancement and weakening in binding affinity. The specific predicted data is provided in **Supplementary Table S4.**

To complement our computational predictions with a structural perspective, we explored the mutant conformations using AlphaFold 3. Although these models are themselves predictive, the observed structural features aligned with the predicted changes in binding affinity. In the H: I102E mutant **(Fig. 6E)**, the substitution of isoleucine with glutamate at position 102 introduced a shorter hydrogen-bonding network with adjacent residues. The negatively charged side chain of glutamate forms additional electrostatic interactions with nearby polar residues, compensating for the loss of hydrophobic contacts originally contributed by isoleucine. This rearrangement potentially enhances local structural rigidity and stabilizes the binding interface. Conversely, the H: D107H **(Fig. 6F)** mutation replaced a negatively charged aspartate with histidine, disrupting a key hydrogen bond and potentially weakening the interaction surface. These in silico observations provide a plausible structural rationale for the predicted effects of mutations on binding affinity, offering mechanistic hypotheses that warrant future experimental validation.

## DISCUSSION

In this study, we present SAMAffinity, a deep learning neural network designed to predict the effects of single and multiple mutations on protein-protein binding affinity changes. Unlike current mainstream graph neural network (GNN) approaches that rely on structural data, SAMAffinity employs a hybrid CNN-Transformer architecture and operates solely on sequence information, enabling efficient and accurate prediction. In performance evaluations, SAMAffinity not only significantly outperforms existing sequence-based baseline methods, but also surpasses advanced structure-based approaches in predictive accuracy on two independent test subsets of the SKEMPI 2.0 dataset.

To enhance the model’s capacity to learn protein-protein interaction patterns, we constructed a multi-dimensional feature representation framework. Firstly, we integrated physicochemical properties along with evolutionary profiles, including BLOSUM62 and PSSM, to comprehensively capture local microenvironmental changes. Secondly, we incorporated sequence embeddings from multiple protein language models (ESM-2, ProtT5 and AntiBERTy), which provide complementary contextual information learned at different semantic levels. Ablation results confirmed that each of these feature types contributes significantly to the overall performance of the model.

To demonstrate the generalizability and practical utility of the model, we applied SAMAffinity to a case study of SARS-CoV-2 and further explored amino acid optimization of the CDR-H3 region of the CR3022 antibody based on SAMAffinity’s predictions. Structural modeling using AlphaFold 3 provided interpretations consistent with the predicted affinity trends, offering computational validation at the molecular scale. Reliable ΔΔG prediction not only facilitates rational protein optimization but also serves as a key metric for evaluating the effectiveness of protein design strategies. Currently, the field of protein sequence design lacks robust methodologies for quantitatively assessing the functional improvement of designed sequences. Accurate ΔΔG prediction can offer physically grounded energetic interpretations for designed variants, shows the potential to fill this critical methodological gap.

In addition, a comprehensive analysis of the experimental results revealed two key limitations that underscore the need for further improvement. A substantial portion of the training data, approximately 40%, is derived from alanine scanning experiments, which may lead the model to develop a bias toward alanine mutation patterns and result in an inherent positive shift in its predictions. Moreover, when predicting mutation effects within the same wild-type complex, the sequence feature tensors often show only minor variations due to single-point substitutions. This can cause the predicted ΔΔG values to cluster within a narrow numerical range, limiting the model’s capacity to resolve subtle changes in binding affinity. These observations provide directions for future methodological enhancements.

## METHODS

### Dataset

To ensure a fair and rigorous comparison with existing methods, our study utilized two widely-adopted public benchmarks, SKEMPI 2.0^42^ and AB-Bind^41^. All samples within these datasets have been manually curated and are accompanied by structural data of the corresponding wild-type complexes. In both datasets, the changes in binding affinity is defined as^40^:

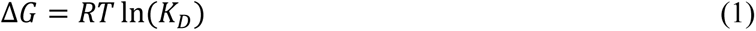

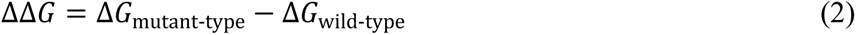

where, 𝑅 = 1.985 × 10^−3^ 𝑘𝑐𝑎𝑙 · 𝐾^−1^ · 𝑚𝑜𝑙^−1^ represents the ideal gas constant, 𝑇 represents the dissociation temperature (in Kelvin), and 𝐾_𝐷_ denotes the equilibrium dissociation constant of the protein complex.

The SKEMPI 2.0 dataset contains 7,085 mutation samples, encompassing various types of protein-protein interactions such as antibody-antigen and protease-inhibitor complexes. Xiong et al.^39^ extracted a non-redundant subset, S1131, from SKEMPI 2.0, which includes only single-point mutations and consists entirely of dimeric complexes. Rodrigues et al.^40^ derived a larger single-point mutation dataset, S4169, from SKEMPI 2.0, comprising 4,169 samples across 319 complexes. AB-Bind is a dataset composed exclusively of antibody-antigen interactions, containing a total of 1,101 samples, and is also referred to as the M1101 dataset. It includes both single-point and multiple-point mutation samples. Following the approach of Wang et al.^18^, a subset designated S645 containing only single-point mutations was extracted from AB-Bind.

### Featurization and model input

#### Physicochemical properties

The mechanism by which amino acid mutations influence binding affinity primarily arises from changes in the chemical properties and functional characteristics of side chains. To systematically characterize the physicochemical changes in the local microenvironment of mutation sites, we selected features such as polarizability, hydrophobicity^35^, secondary-structure, and relative accessible surface area (RASA)^36^. Among these, the RASA was predicted using the NetSurfP-3.0 online server. The computational procedures for all other features are provided in **Supplementary Table S1.**

#### Evolutionary profiles

During biological evolution, amino acid substitutions exhibit pronounced conservation patterns. To quantitatively assess substitution tendencies, we employed the BLOSUM62^37^ and position-specific scoring matrix (PSSM)^38^. BLOSUM62, as a global substitution matrix, reflects amino acid replacement patterns observed in distant evolutionary relationships and does not account for specific protein positions. In contrast, PSSM captures evolutionary conservation within specific protein families through multiple sequence alignment, thereby representing site-specific tolerance to amino acid mutations. PSSM was generated using the PSI-BLAST program from BLAST 2.12.0, with the Swiss-Prot database as the search source. Since NetSurfP-3.0 and PSI-BLAST are not applicable to short peptide sequences, RASA and PSSM features were therefore excluded for the corresponding samples in the S4169 dataset.

#### Protein language model embeddings

In addition to the aforementioned handcrafted features, we also employed ESM-2^28^ and ProtT5^27^ to extract general sequence features. ESM-2 predicts randomly masked amino acid tokens by leveraging contextual information, thereby effectively capturing residue-level contextual dependencies. In contrast, ProtT5, benefiting from its encoder-decoder architecture that predicts masked residue spans token by token, is better suited for sequence generation tasks. Notably, given the unique characteristics of antibody-antigen interactions, we replaced ProtT5 with AntiBERTy^26^ during training on the AB-Bind dataset to generate antibody-specific embeddings. AntiBERTy is a BERT-based model trained on a non-redundant dataset of 558 million antibody sequences. Compared to general protein language models, AntiBERTy is better equipped to capture affinity maturation trajectories existed in antibody sequence data.

The specific shapes and parameters of all the aforementioned features are provided in the **Supplementary Table S2**.

### Network architecture

The overall architecture of the network is illustrated in **Figure 1C**. To more accurately simulate the binding process of protein-protein complexes, the extracted features are divided into four components: receptor-wild-type, receptor-mutant-type, ligand-wild-type, and ligand-mutant-type. These components are then fed into two pairs of encoder-decoder modules with non-shared parameters.

#### Encoder design

The encoder consists of an AFC module. Each group of features, represented as 𝑥_𝑖_ ∈ 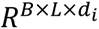, where 𝑖 denotes the 𝑖-th feature, is first passed through independent linear layers for weight learning and dimension alignment. Then the weighted feature tensors are concatenated along the feature dimension to form a unified representation 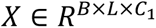, a 1D residual convolutional module is employed to extract local contextual features for each component, outputting high-dimension local convolutional representations 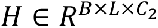:

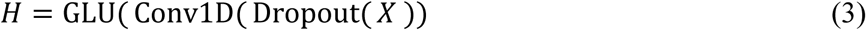

where, 𝐻 comprises 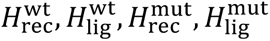.

#### Decoder design

The decoder is composed of two core modules: MSI and BIA. Although MSI module and BIA module share an identical Transformer-based cross-attention architecture, they are functionally distinguished by their inputs and learning objectives. Specifically, MSI focuses on local environmental perturbations around the mutation site, while BIA captures cooperative mutation effects across the receptor-ligand interface. The design of these blocks is defined as follows:

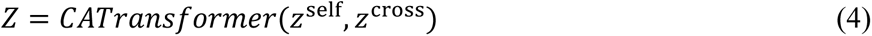

For the MSI module, which is employed to adaptively capture the significant differences between the wild-type and mutant-type sequence representations, is defined as follows:

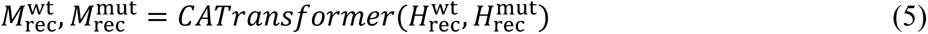

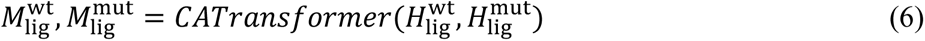

where, these two blocks share the same architecture as defined in **Eq. 4**, but their parameters are not shared.

For the BIA module, which is employed to model the propagation of mutation effects at the binding interface and systematically capturing cooperative change patterns among interface residues, is defined as follows:

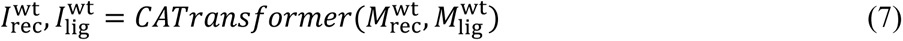

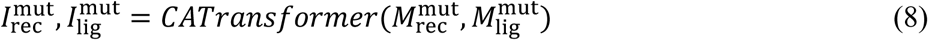

where, these two blocks share the same architecture as defined in **Eq. 4**, but their parameters are not shared. We adopted this design because the impact of a mutation goes beyond the amino acid change itself, depending critically on the role of the mutation site within the broader sequence context. Proteins are not linear residue chains but structured folds, where each residue’s function is shaped by its surrounding environment.

#### Output

Based on the residue-level representations 𝐼 from the IAM, we aggregate them into sequence-level representations *Z* via L2-norm weighted pooling. The complex-level representations for the wild-type (*Z* ^wt^) and mutant-type (Z ^mut^), are then concatenated. An explicit differential term (*Z* ^wt^ - Z ^mut^) is incorporated to physically fit the binding affinity changes. Finally, fully connected layers are used to predict the ΔΔG values for both the forward (wild to mutant, ΔΔG_Forward_) and reverse (mutant to wild, ΔΔG_Reverse_) processes, both of which contribute to the model’s loss function during training.

### Training strategy

To mitigate the impact of class imbalance between positive and negative samples on model prediction performance, we adopted the following loss function inspired by Zhou et al.^46^, replacing the conventional mean squared error loss:

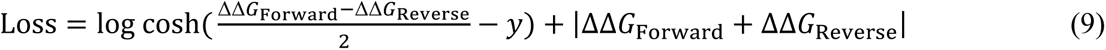

where, *y* denotes the ground truth label. Theoretically, the model’s predictions should satisfy the following symmetry constraints:

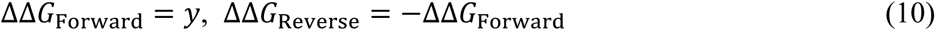

This loss function is designed to enhance the model’s ability to learn the symmetry between forward and reverse mutations, thereby alleviating inherent biases within the dataset. Specifically, all models were trained for 150 epochs, with the learning rate decaying every 5 epochs. RAdam was selected as the optimizer to balance training stability and convergence efficiency, with the hyperparameters set to (β_1_, β_2_) = (0.9, 0.999) and ε = 1×10⁻⁸. Model checkpoints were selected based on the Pearson correlation coefficients. Regarding hardware configuration, a single round of 10-fold cross-validation required approximately 22 hours of training on an NVIDIA Tesla V100S-PCIE-32GB GPU. Other hyperparameters, such as batch size and initial learning rate are detailed in the **Supplementary Table S5.**

## ACKNOWLEDGEMENTS

We thank members of the G. Z. lab for discussion and feedback; R. Zheng for helping with dataset preparations. Computational resources were provided by the College of Information Engineering at Zhejiang University of Technology. This work was supported by the National Key R&D Program of China [2022ZD0115103, G.Z.], the National Nature Science Foundation of China [62173304, G.Z.; 62573386, G.Z.], the “Pioneer” and “Leading Goose” R&D Program of Zhejiang [2025C01190, G.Z.], and the Zhejiang Province High-level Talent Special Support Program [2023R5248, G.Z.].

## DATA AVAILABILITY

SAMAffinity is available for download via GitHub at https://github.com/iobio-zjut/SAMAffinity.

## SUPPLEMENTARY NOTES

### Supplementary Note S1. Steiger’s Z-test for comparing dependent correlations

To assess whether our model achieved a statistically higher correlation with the ground truth compared to the baseline model, we applied Steiger’s Z-test, which is designed to compare two dependent correlations sharing one common variable.

#### (1) Problem definition

Let

- 𝑌: the ground truth (real distribution),
- 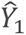: predictions from our model,
- 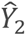: predictions from the baseline model.

We aim to test whether the correlations:

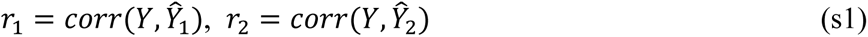

are significantly different:

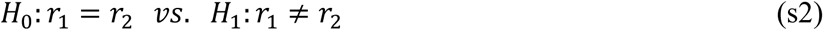

Since both correlations involve the same variable 𝑌, they are statistically dependent. The correlation between the two prediction sets is denoted as 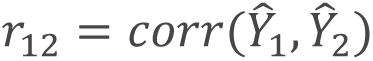

#### (2) Fisher’s z-transformation

Each correlation coefficient is first transformed using the Fisher’s z-transform to approximate normality:

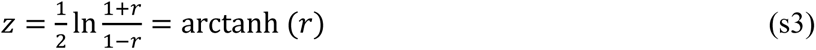

#### (3) Steiger’s Z-test

The Steiger’s Z-test for comparing the two dependent correlations is computed as:

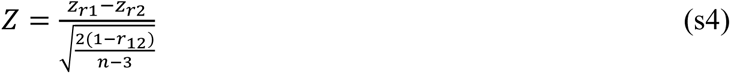

where, *n* represents the number of samples.

#### (4) *P*-value calculation

The two-tailed *P*-value is given by:

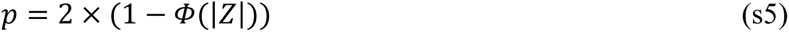

where, 𝛷(·) denotes the cumulative distribution function of the standard normal distribution.

### Supplementary Note S2. Cross validation evaluation strategy

For mutation-level evaluation, we employed 10-fold cross-validation to assess model’s performance. Specifically, the K-Fold function from the scikit-learn library was used for data partitioning, with a fixed random seed to ensure reproducibility. To prevent data leakage during prediction, we adopted the following evaluation strategy: each fold’s model was trained independently and used to predict its corresponding validation set. The predictions from all validation sets were then aggregated to compute the overall performance metrics. Compared to simply averaging the performance of 10 individual models, this approach more accurately reflects the model’s predictive consistency on the global dataset, reduces evaluation bias caused by random partitioning, and aligns more closely with practical application scenarios.

**Supplementary Fig. 1.**
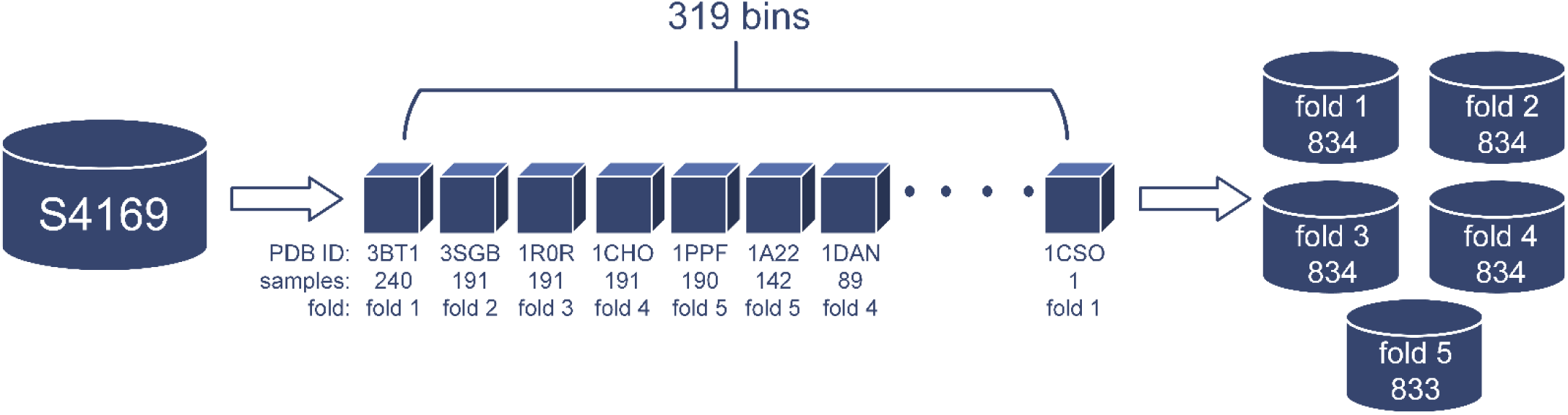
Schematic illustration of the complex-level data splitting procedure.

For complex-level evaluation, we followed the 5-fold cross-validation protocol established by the benchmark method MpbPPI^20^. The detailed implementation steps are as follows: (i) The dataset is partitioned into bins at the complex level. Taking S4169 as an example, which consists of 319 complexes, it is divided into 319 bins accordingly. (ii) The bins are then sorted in descending order based on the number of samples they contain. (iii) The sorted bins are assigned to 5 folds using a greedy algorithm to ensure that the number of samples in each fold is approximately balanced. This partitioning strategy strictly ensures that mutation samples from the same complex do not appear simultaneously in both the training and validation sets, thereby preventing the model from encountering data patterns in training that are highly similar to those in the validation set. During performance evaluation, the same assessment strategy used for mutation-level validation is applied.

## SUPPLEMENTARY TABLES

**Supplementary Table S1.** Amino acid properties. Table S1 lists the values of seven physicochemical properties corresponding to the 20 amino acids^35^.

| Amino acid | Steric parameter | Polarizability | Volume | Hydrophobicity | Isoelectric pt | Helix prob | Sheet prob |
| --- | --- | --- | --- | --- | --- | --- | --- |
| ALA | 1.28 | 0.05 | 1 | 0.31 | 6.11 | 0.42 | 0.23 |
| GLY | 0 | 0 | 0 | 0 | 6.07 | 0.13 | 0.15 |
| VAL | 3.67 | 0.14 | 3 | 1.22 | 6.02 | 0.27 | 0.49 |
| LEU | 2.59 | 0.19 | 4 | 1.7 | 6.04 | 0.39 | 0.31 |
| ILE | 4.19 | 0.19 | 4 | 1.8 | 6.04 | 0.3 | 0.45 |
| PHE | 2.94 | 0.29 | 5.89 | 1.79 | 5.67 | 0.3 | 0.38 |
| TYR | 2.94 | 0.3 | 6.47 | 0.96 | 5.66 | 0.25 | 0.41 |
| TRP | 3.21 | 0.41 | 8.08 | 2.25 | 5.94 | 0.32 | 0.42 |
| THR | 3.03 | 0.11 | 2.6 | 0.26 | 5.6 | 0.21 | 0.36 |
| SER | 1.31 | 0.06 | 1.6 | -0.04 | 5.7 | 0.2 | 0.28 |
| ARG | 2.34 | 0.29 | 6.13 | -1.01 | 10.74 | 0.36 | 0.25 |
| LYS | 1.89 | 0.22 | 4.77 | -0.99 | 9.99 | 0.32 | 0.27 |
| HIS | 2.99 | 0.23 | 4.66 | 0.13 | 7.69 | 0.27 | 0.3 |
| ASP | 1.6 | 0.11 | 2.78 | -0.77 | 2.95 | 0.25 | 0.2 |
| GLU | 1.56 | 0.15 | 3.78 | -0.64 | 3.09 | 0.42 | 0.21 |
| ASN | 1.6 | 0.13 | 2.95 | -0.6 | 6.52 | 0.21 | 0.22 |
| GLN | 1.56 | 0.18 | 3.95 | -0.22 | 5.65 | 0.36 | 0.25 |
| MET | 2.35 | 0.22 | 4.43 | 1.23 | 5.71 | 0.38 | 0.32 |
| PRO | 2.67 | 0 | 2.72 | 0.72 | 6.8 | 0.13 | 0.34 |
| CYS | 1.77 | 0.13 | 2.43 | 1.54 | 6.35 | 0.17 | 0.41 |

**Supplementary Table S2.** Shape of features.

| Hyperparameters | Shape |
| --- | --- |
| Physicochemical properties | L*7 |
| Secondary structure | L*3 |
| Relative accessible surface area | L*1 |
| BLOSUM62 | L*23 |
| PSSM | L*20 |
| ESM-2 embedding | L*1280 |
| ProtT5 embedding | L*1024 |
| AntiBERTy embedding | L*512 |

**Supplementary Table S3.** SARS-COV-2 test set. Table S3 lists all sample points used in the SARS-COV-2 test set^33, 34^, including antibody type, strain type, mutation information, and experimentally measured ΔΔG.

| Antibody | Antigen | Mutation_info | ddG |
| --- | --- | --- | --- |
| CR3022 | SARS-COV-2 Wild | R: A372T | 0.014 |
| CR3022 | SARS-COV-2 Wild | R: P384A | 2.302 |
| CR3022 | SARS-COV-2 Wild | R: T430M | 0.037 |
| CR3022 | SARS-COV-2 Wild | R: H519N | 0.069 |
| C105 | SARS-COV-2 Wild | R: R346S | 0.212 |
| C105 | SARS-COV-2 Wild | R: N439K | 0.181 |
| C105 | SARS-COV-2 Wild | R: N440K | 0.149 |
| C105 | SARS-COV-2 Wild | R: A475V | 1.647 |
| C105 | SARS-COV-2 Wild | R: V483A | 0.041 |
| C105 | SARS-COV-2 Wild | R: E484K | 0.181 |
| C105 | SARS-COV-2 Wild | R: Q493R | -0.502 |
| C144 | SARS-COV-2 Wild | R: R346S | 0.629 |
| C144 | SARS-COV-2 Wild | R: N439K | 0.283 |
| C144 | SARS-COV-2 Wild | R: N440K | 0.474 |
| C144 | SARS-COV-2 Wild | R: A475V | 1.023 |
| C144 | SARS-COV-2 Wild | R: V483A | 0.341 |
| C002 | SARS-COV-2 Wild | R: R346S | 0.532 |
| C002 | SARS-COV-2 Wild | R: N439K | 0.222 |
| C002 | SARS-COV-2 Wild | R: N440K | 0.463 |
| C002 | SARS-COV-2 Wild | R: A475V | 0.614 |
| C002 | SARS-COV-2 Wild | R: V483A | 0 |
| C002 | SARS-COV-2 Wild | R: Q493R | 2.368 |
| C121 | SARS-COV-2 Wild | R: R346S | 0.279 |
| C121 | SARS-COV-2 Wild | R: N439K | 0.279 |
| C121 | SARS-COV-2 Wild | R: N440K | 0.108 |
| C121 | SARS-COV-2 Wild | R: A475V | 0.200 |
| C121 | SARS-COV-2 Wild | R: V483A | 0.279 |
| C121 | SARS-COV-2 Wild | R: Q493R | 3.204 |
| C119 | SARS-COV-2 Wild | R: R346S | 0.468 |
| C119 | SARS-COV-2 Wild | R: N439K | 0.411 |
| C119 | SARS-COV-2 Wild | R: N440K | 0.440 |
| C119 | SARS-COV-2 Wild | R: A475V | 0.411 |
| C119 | SARS-COV-2 Wild | R: V483A | 0.468 |
| C119 | SARS-COV-2 Wild | R: E484K | 1.31 |
| C119 | SARS-COV-2 Wild | R: Q493R | 0.411 |
| C104 | SARS-COV-2 Wild | R: R346S | 0.967 |
| C104 | SARS-COV-2 Wild | R: N439K | 0.139 |
| C104 | SARS-COV-2 Wild | R: N440K | 1.03 |
| C104 | SARS-COV-2 Wild | R: A475V | 1.112 |
| C104 | SARS-COV-2 Wild | R: V483A | 0.765 |
| C104 | SARS-COV-2 Wild | R: Q493R | -0.066 |
| C135 | SARS-COV-2 Wild | R: N439K | 1.079 |
| C135 | SARS-COV-2 Wild | R: Q493R | 0.459 |
| C110 | SARS-COV-2 Wild | R: R346S | 2.347 |
| C110 | SARS-COV-2 Wild | R: N439K | 1.078 |
| C110 | SARS-COV-2 Wild | R: N440K | 0.255 |
| C110 | SARS-COV-2 Wild | R: A475V | 0.496 |
| C110 | SARS-COV-2 Wild | R: E484K | 1.799 |
| C110 | SARS-COV-2 Wild | R: Q493R | 1.318 |
| SARS VHH-72 | SARS-COV-2 Wild | L: S56M | -0.830 |
| SARS VHH-72 | SARS-COV-2 Wild | L: L97W | -1.069 |
| SARS VHH-72 | SARS-COV-2 Wild | L: T99V | -1.698 |
| SARS VHH-72 | SARS-COV-2 Wild | L: S56M, L: L97W | -2.015 |
| SARS VHH-72 | SARS-COV-2 Wild | L: S56M, L: T99V | -2.082 |
| SARS VHH-72 | SARS-COV-2 Wild | L: L97W, L: T99V | -2.165 |
| SARS VHH-72 | SARS-COV-2 Wild | L: S56M, L: L97W, L: T99V | -2.331 |
| SARS VHH-72 | SARS-COV-2 B.1.351 | L: S56M | -0.247 |
| SARS VHH-72 | SARS-COV-2 B.1.351 | L: L97W | -0.358 |
| SARS VHH-72 | SARS-COV-2 B.1.351 | L: T99V | -0.391 |
| SARS VHH-72 | SARS-COV-2 B.1.351 | L: S56M, L: L97W | -0.699 |
| SARS VHH-72 | SARS-COV-2 B.1.351 | L: S56M, L: T99V | -1.106 |
| SARS VHH-72 | SARS-COV-2 B.1.351 | L: L97W, L: T99V | -1.123 |
| SARS VHH-72 | SARS-COV-2 B.1.351 | L: S56M, L: L97W, L: T99V | -1.257 |
| SARS VHH-72 | SARS-COV-2 B.1.617.2 | L: S56M | -0.495 |
| SARS VHH-72 | SARS-COV-2 B.1.617.2 | L: L97W | -0.506 |
| SARS VHH-72 | SARS-COV-2 B.1.617.2 | L: T99V | -0.622 |
| SARS VHH-72 | SARS-COV-2 B.1.617.2 | L: S56M, L: L97W | -0.731 |
| SARS VHH-72 | SARS-COV-2 B.1.617.2 | L: S56M, L: T99V | -0.799 |
| SARS VHH-72 | SARS-COV-2 B.1.617.2 | L: L97W, L: T99V | -0.981 |
| SARS VHH-72 | SARS-COV-2 B.1.617.2 | L: S56M, L: L97W, L: T99V | -0.910 |

**Supplementary Table S4.** Specific prediction data of CR3022 CDR-H3 region.

| Mutant type | G99 | S100 | G101 | I102 | S103 | T104 | P105 | M106 | D107 | V108 |
| --- | --- | --- | --- | --- | --- | --- | --- | --- | --- | --- |
| G | / | -0.98 | / | 1.55 | -1.28 | 0.33 | 0.52 | 0.41 | 1.73 | -0.68 |
| A | -0.06 | -0.70 | -0.87 | 3.49 | -1.54 | -0.04 | 0.17 | -1.06 | -0.37 | -0.30 |
| V | 0.79 | -1.03 | 0.05 | -0.43 | -0.43 | -0.44 | 0.79 | 2.06 | 0.29 | / |
| L | 0.62 | 0.83 | -0.96 | 1.04 | 0.24 | -0.17 | 0.66 | -1.29 | 2.46 | -1.67 |
| I | 1.04 | -1.06 | 0.07 | / | -0.32 | -0.83 | 0.82 | 0.83 | 0.41 | -1.43 |
| P | -0.05 | -0.54 | 0.57 | -0.27 | -0.15 | -1.43 | / | 0.17 | 1.07 | -1.32 |
| F | 0.94 | 0.05 | -0.17 | -0.67 | 2.11 | -0.76 | -0.21 | -1.27 | -0.06 | -0.97 |
| Y | 0.47 | 0.53 | -0.33 | 0.14 | -0.09 | -0.89 | -0.35 | -0.57 | 1.62 | -1.51 |
| W | 0.54 | 0.15 | -0.14 | -0.05 | -0.23 | -0.80 | -0.24 | 0.39 | -0.39 | -1.39 |
| S | -0.74 | / | -0.26 | 0.46 | / | 0.42 | 0.05 | 0.47 | -0.75 | -0.54 |
| T | -0.06 | -1.21 | 0.04 | 1.32 | -0.59 | / | -0.12 | -0.05 | -0.23 | -0.58 |
| C | 0.41 | -0.25 | -0.11 | 1.48 | -1.57 | -1.34 | -0.22 | 1.59 | 0.88 | -1.33 |
| M | 0.57 | -0.02 | -0.15 | -0.08 | -0.66 | -0.22 | 0.34 | / | 3.87 | 0.70 |
| N | -0.80 | -0.38 | -0.16 | 0.44 | 0.32 | 0.27 | 0.98 | 0.72 | -1.18 | 2.30 |
| Q | -1.26 | -0.27 | 0.19 | 1.84 | -0.13 | -1.10 | 0.28 | 0.90 | -1.39 | 0.48 |
| D | -1.19 | -0.45 | -1.00 | 0.68 | -0.58 | 1.23 | 0.17 | 2.53 | / | -0.31 |
| E | -1.14 | -0.16 | 0.03 | 3.46 | -1.28 | -0.07 | -0.09 | 0.34 | -0.40 | 0.18 |
| K | -1.21 | 0.89 | 0.17 | 1.57 | -0.17 | -0.54 | -0.16 | -0.60 | -1.43 | -0.51 |
| R | -1.32 | -0.37 | -0.16 | 2.30 | 0.28 | -0.49 | 0.03 | 0.06 | -1.44 | 1.41 |
| H | -1.27 | -0.88 | 0.72 | 0.24 | -0.10 | -1.09 | 0.12 | 1.85 | -1.82 | 0.36 |

**Supplementary Table S5.**
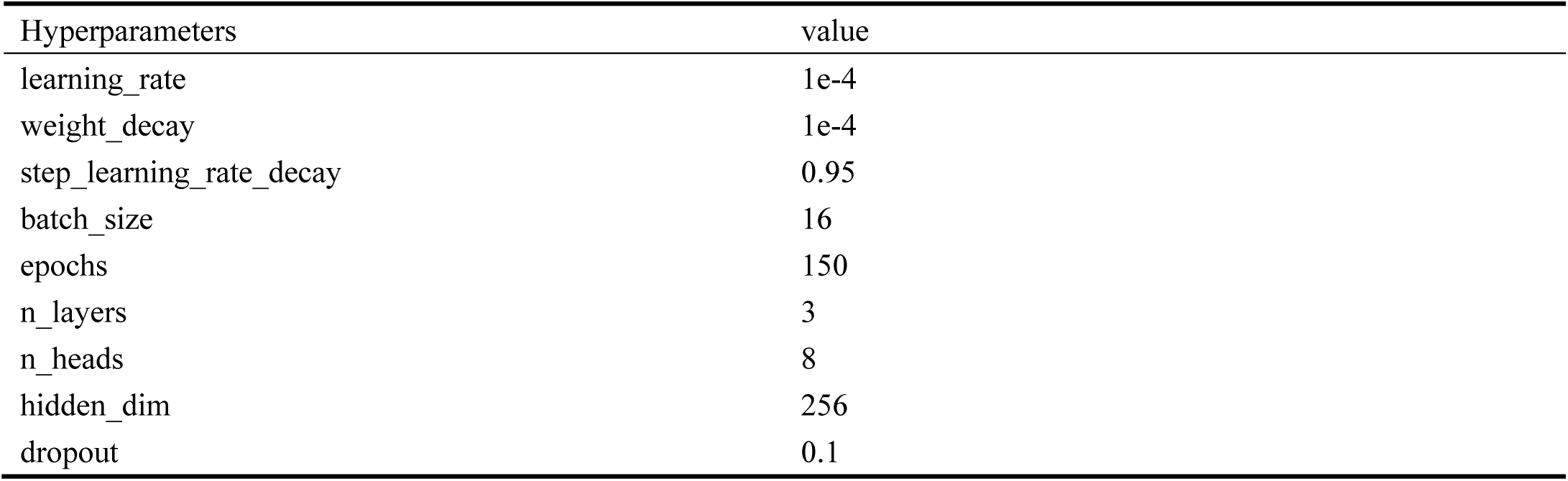
Supplementary hyperparameters.

| Hyperparameters | value |
| --- | --- |
| learning_rate | 1e-4 |
| weight_decay | 1e-4 |
| step_learning_rate_decay | 0.95 |
| batch_size | 16 |
| epochs | 150 |
| n_layers | 3 |
| n_heads | 8 |
| hidden_dim | 256 |
| dropout | 0.1 |

## SUPPLEMENTARY ALGORITHMS

**Supplementary Algorithm S1.**
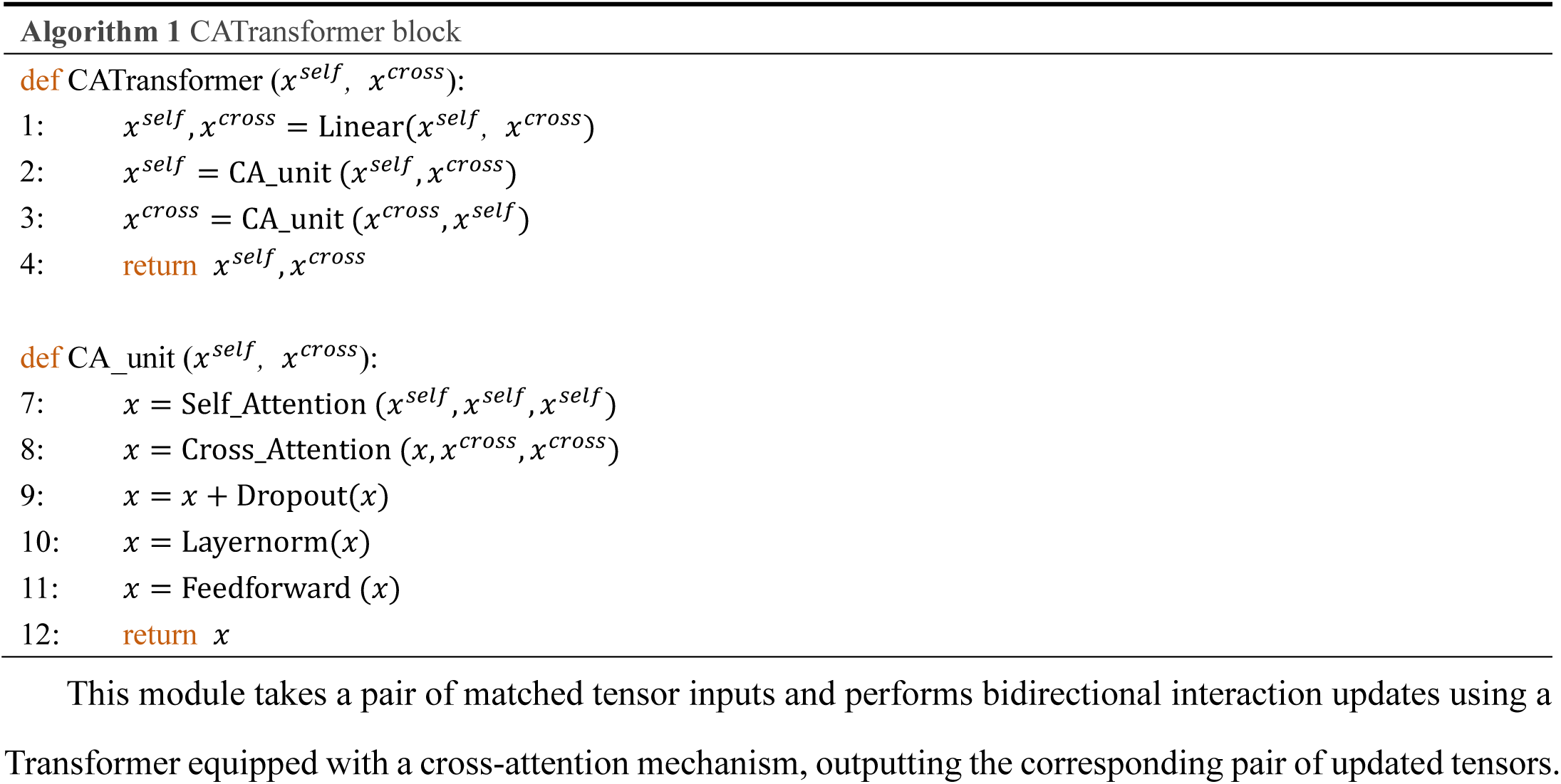
CATransformer block design.

## REFERENCES

1. Zhou, Z.; Yin, Y.; Han, H.; Jia, Y.; Koh, J. H.; Kong, A. W.-K.; Mu, Y., ProAffinity-GNN: A Novel Approach to Structure-Based Protein-Protein Binding Affinity Prediction via a Curated Data Set and Graph Neural Networks. Journal of Chemical Information and Modeling 2024, 64, 8796–8808.

2. Dehouck, Y.; Kwasigroch, J. M.; Rooman, M.; Gilis, D., BeAtMuSiC: prediction of changes in protein-protein binding affinity on mutations. Nucleic Acids Research 2013, 41, W333–W339.

3. Pires, D. E. V.; Ascher, D. B., mCSM-AB: a web server for predicting antibody-antigen affinity changes upon mutation with graph-based signatures. Nucleic Acids Research 2016, 44, W469–W473.

4. Li, M.; Petukh, M.; Alexov, E.; Panchenko, A. R., Predicting the Impact of Missense Mutations on Protein-Protein Binding Affinity. Journal of Chemical Theory and Computation 2014, 10, 1770–1780.

5. Starr, T. N.; Greaney, A. J.; Hilton, S. K.; Ellis, D.; Crawford, K. H. D.; Dingens, A. S.; Navarro, M. J.; Bowen, J. E.; Tortorici, M. A.; Walls, A. C.; King, N. P.; Veesler, D.; Bloom, J. D., Deep Mutational Scanning of SARS-CoV-2 Receptor Binding Domain Reveals Constraints on Folding and ACE2 Binding. Cell 2020, 182, 1295-+.

6. ProstaNet: A Novel Geometric Vector Perceptrons-Graph Neural Network Algorithm for Protein Stability Prediction in Single- and Multiple-Point Mutations with Experimental Validation. Research 2025, 2025.

7. French, D. L.; Laskov, R.; Scharff, M. D., The role of somatic hypermutation in the generation of antibody diversity. *Science (New York*, N.Y.) 1989, 244, 1152–1157.

8. Mesin, L.; Ersching, J.; Victora, G. D., Germinal Center B Cell Dynamics. Immunity 2016, 45, 471–482.

9. Di Nola, J. M.; Neuberger, M. S., Molecular mechanisms of antibody somatic hypermutation. Annual Review of Biochemistry 2007, 76, 1–22.

10. Willander, M.; Al-Hilli, S. Analysis of Biomolecules Using Surface Plasmons. In Micro and Nano Technologies in Bioanalysis: Methods and Protocols, Lee, J. W.; Foote, R. S., Eds.; 2009; Vol. 544, pp 201–229.

11. Ladbury, J. E.; Chowdhry, B. Z., Sensing the heat: The application of isothermal titration calorimetry to thermodynamic studies of biomolecular interactions. Chemistry and Biology (London) 1996, 3, 791–801.

12. Tsishyn, M.; Pucci, F.; Rooman, M., Quantification of biases in predictions of protein-protein binding affinity changes upon mutations. Briefings in Bioinformatics 2024, 25.

13. Schymkowitz, J.; Borg, J.; Stricher, F.; Nys, R.; Rousseau, F.; Serrano, L., The FoldX web server: an online force field. Nucleic Acids Research 2005, 33, W382–W388.

14. Alford, R. F.; Leaver-Fay, A.; Jeliazkov, J. R.; O’Meara, M. J.; DiMaio, F. P.; Park, H.; Shapovalov, M. V.; Renfrew, P. D.; Mulligan, V. K.; Kappel, K.; Labonte, J. W.; Pacella, M. S.; Bonneau, R.; Bradley, P.; Dunbrack, R. L., Jr.; Das, R.; Baker, D.; Kuhlman, B.; Kortemme, T.; Gray, J. J., The Rosetta All-Atom Energy Function for Macromolecular Modeling and Design. Journal of Chemical Theory and Computation 2017, 13, 3031–3048.

15. Barlow, K. A.; Conchuir, S. O.; Thompson, S.; Suresh, P.; Lucas, J. E.; Heinonen, M.; Kortemme, T., Flex ddG: Rosetta Ensemble-Based Estimation of Changes in Protein-Protein Binding Affinity upon Mutation. Journal of Physical Chemistry B 2018, 122, 5389–5399.

16. Marabotti, A.; Scafuri, B.; Facchiano, A., Predicting the stability of mutant proteins by computational approaches: an overview. Briefings in Bioinformatics 2021, 22.

17. Zhang, N.; Chen, Y.; Lu, H.; Zhao, F.; Alvarez, R. V.; Goncearenco, A.; Panchenko, A. R.; Li, M., MutaBind2: Predicting the Impacts of Single and Multiple Mutations on Protein-Protein Interactions. Iscience 2020, 23.

18. Wang, M.; Cang, Z.; Wei, G.-W., A topology-based network tree for the prediction of protein-protein binding affinity changes following mutation. Nature Machine Intelligence 2020, 2, 116–123.

19. Liu, X.; Luo, Y.; Li, P.; Song, S.; Peng, J., Deep geometric representations for modeling effects of mutations on protein-protein binding affinity. Plos Computational Biology 2021, 17.

20. Yue, Y.; Li, S.; Wang, L.; Liu, H.; Tong, H. H. Y.; He, S., MpbPPI: a multi-task pre-training-based equivariant approach for the prediction of the effect of amino acid mutations on protein-protein interactions. Briefings in Bioinformatics 2023, 24.

21. Zhou, Y.; Myung, Y.; Rodrigues, C. H. M.; Ascher, D. B., DDMut-PPI: predicting effects of mutations on protein-protein interactions using graph-based deep learning. Nucleic Acids Research 2024, 52, W207–W214.

22. Cai, H.; Zhang, Z.; Wang, M.; Zhong, B.; Li, Q.; Zhong, Y.; Wu, Y.; Ying, T.; Tang, J., Pretrainable geometric graph neural network for antibody affinity maturation. Nature Communications 2024, 15.

23. Scarselli, F.; Gori, M.; Tsoi, A. C.; Hagenbuchner, M.; Monfardini, G., The Graph Neural Network Model. IEEE Transactions on Neural Networks 2009, 20, 61–80.

24. Vaswani, A.; Shazeer, N.; Parmar, N.; Uszkoreit, J.; Jones, L.; Gomez, A. N.; Kaiser, Ł.; Polosukhin, I., Attention is all you need. Advances in neural information processing systems 2017, 30.

25. Siebenmorgen, T.; Zacharias, M., Computational prediction of protein-protein binding affinities. Wiley Interdisciplinary Reviews-Computational Molecular Science 2020, 10.

26. Ruffolo, J. A.; Gray, J. J.; Sulam, J., Deciphering antibody affinity maturation with language models and weakly supervised learning. arXiv preprint arXiv:2112.07782 2021.

27. Elnaggar, A.; Heinzinger, M.; Dallago, C.; Rehawi, G.; Wang, Y.; Jones, L.; Gibbs, T.; Feher, T.; Angerer, C.; Steinegger, M.; Bhowmik, D.; Rost, B., ProtTrans: Toward Understanding the Language of Life Through Self-Supervised Learning. Ieee Transactions on Pattern Analysis and Machine Intelligence 2022, 44, 7112–7127.

28. Lin, Z.; Akin, H.; Rao, R.; Hie, B.; Zhu, Z.; Lu, W.; Smetanin, N.; Verkuil, R.; Kabeli, O.; Shmueli, Y.; Costa, A. d. S.; Fazel-Zarandi, M.; Sercu, T.; Candido, S.; Rives, A., Evolutionary-scale prediction of atomic-level protein structure with a language model. Science 2023, 379, 1123–1130.

29. Rao, R.; Bhattacharya, N.; Thomas, N.; Duan, Y.; Chen, X.; Canny, J.; Abbeel, P.; Song, Y. S. Evaluating Protein Transfer Learning with TAPE. In 33rd Conference on Neural Information Processing Systems (NeurIPS), Vancouver, CANADA, 2019 Dec 08-14.

30. Zhang, C.; Sun, Y.; Hu, P., An interpretable deep geometric learning model to predict the effects of mutations on protein–protein interactions using large-scale protein language model. Journal of Cheminformatics 2025, 17, 35.

31. Jin, R.; Ye, Q.; Wang, J.; Cao, Z.; Jiang, D.; Wang, T.; Kang, Y.; Xu, W.; Hsieh, C.-Y.; Hou, T., AttABseq: an attention-based deep learning prediction method for antigen-antibody binding affinity changes based on protein sequences. Briefings in Bioinformatics 2024, 25.

32. Xia, Y.; Wang, Z.; Huang, F.; Xiong, Z.; Wang, Y.; Qiu, M.; Zhang, W., DeepInterAware: Deep Interaction Interface-Aware Network for Improving Antigen-Antibody Interaction Prediction from Sequence Data. Advanced Science 2025, 12.

33. Barnes, C. O.; Jette, C. A.; Abernathy, M. E.; Dam, K.-M. A.; Esswein, S. R.; Gristick, H. B.; Malyutin, A. G.; Sharaf, N. G.; Huey-Tubman, K. E.; Lee, Y. E.; Robbiani, D. F.; Nussenzweig, M. C.; West, A. P., Jr.; Bjorkman, P. J., SARS-CoV-2 neutralizing antibody structures inform therapeutic strategies. Nature 2020, 588, 682.

34. Sulea, T.; Baardsnes, J.; Stuible, M.; Rohani, N.; Tran, A.; Parat, M.; Donates, Y. C.; Duchesne, M.; Plante, P.; Kour, G.; Durocher, Y., Structure-based dual affinity optimization of a SARS-CoV-1/2 cross-reactive single-domain antibody. Plos One 2022, 17.

35. Meiler, J.; Müller, M.; Zeidler, A.; Schmäschke, F., Generation and evaluation of dimension-reduced amino acid parameter representations by artificial neural networks. Journal of Molecular Modeling 2001, 7, 360–369.

36. Hoie, M. H.; Kiehl, E. N.; Petersen, B.; Nielsen, M.; Winther, O.; Nielsen, H.; Hallgren, J.; Marcatili, P., NetSurfP-3.0: accurate and fast prediction of protein structural features by protein language models and deep learning. Nucleic Acids Research 2022, 50, W510–W515.

37. Henikoff, S.; Henikoff, J. G., Amino acid substitution matrices from protein blocks. Proceedings of the National Academy of Sciences of the United States of America 1992, 89, 10915–10919.

38. Altschul, S. F.; Madden, T. L.; Schaeffer, A. A.; Zhang, J.; Zhang, Z.; Miller, W.; Lipman, D. J., Gapped BLAST and PSI - BLAST: A new generation of protein database search programs. Nucleic Acids Research 1997, 25, 3389–3402.

39. Xiong, P.; Zhang, C.; Zheng, W.; Zhang, Y., BindProfX: Assessing Mutation-Induced Binding Affinity Change by Protein Interface Profiles with Pseudo-Counts. Journal of Molecular Biology 2017, 429, 426–434.

40. Rodrigues, C. H. M.; Myung, Y.; Pires, D. E. V.; Ascher, D. B., mCSM-PPI2: predicting the effects of mutations on protein-protein interactions. Nucleic Acids Research 2019, 47, W338–W344.

41. Sirin, S.; Apgar, J. R.; Bennett, E. M.; Keating, A. E., AB-Bind: Antibody binding mutational database for computational affinity predictions. Protein Science 2016, 25, 393–409.

42. Jankauskaite, J.; Jimenez-Garcia, B.; Dapkunas, J.; Fernandez-Recio, J.; Moal, I. H., SKEMPI 2.0: an updated benchmark of changes in protein-protein binding energy, kinetics and thermodynamics upon mutation. Bioinformatics 2019, 35, 462–469.

43. Buskirk, A. R.; Green, R., Getting Past Polyproline Pauses. Science 2013, 339, 38–39.

44. Kortemme, T.; Kim, D. E.; Baker, D., Computational alanine scanning of protein-protein interfaces. Science’s STKE : signal transduction knowledge environment 2004, 2004, pl2.

45. Abramson, J.; Adler, J.; Dunger, J.; Evans, R.; Green, T.; Pritzel, A.; Ronneberger, O.; Willmore, L.; Ballard, A. J.; Bambrick, J.; Bodenstein, S. W.; Evans, D. A.; Hung, C.-C.; O’Neill, M.; Reiman, D.; Tunyasuvunakool, K.; Wu, Z.; Zemgulyte, A.; Arvaniti, E.; Beattie, C.; Bertolli, O.; Bridgland, A.; Cherepanov, A.; Congreve, M.; Cowen-Rivers, A. I.; Cowie, A.; Figurnov, M.; Fuchs, F. B.; Gladman, H.; Jain, R.; Khan, Y. A.; Low, C. M. R.; Perlin, K.; Potapenko, A.; Savy, P.; Singh, S.; Stecula, A.; Thillaisundaram, A.; Tong, C.; Yakneen, S.; Zhong, E. D.; Zielinski, M.; Zidek, A.; Bapst, V.; Kohli, P.; Jaderberg, M.; Hassabis, D.; Jumper, J. M., Accurate structure prediction of biomolecular interactions with AlphaFold 3. Nature 2024, 630.

46. Zhou, Y.; Pan, Q.; Pires, D. E. V.; Rodrigues, C. H. M.; Ascher, D. B., DDMut: predicting effects of mutations on protein stability using deep learning. Nucleic Acids Research 2023, 51, W122–W128.

## SUPPLEMENTARY REFERENCES

1. Yue, Y.; Li, S.; Wang, L.; Liu, H.; Tong, H. H. Y.; He, S., MpbPPI: a multi-task pre-training-based equivariant approach for the prediction of the effect of amino acid mutations on protein-protein interactions. Briefings in Bioinformatics 2023, 24.

2. Meiler, J.; Müller, M.; Zeidler, A.; Schmäschke, F., Generation and evaluation of dimension-reduced amino acid parameter representations by artificial neural networks. Journal of Molecular Modeling 2001, 7, 360–369.

3. Barnes, C. O.; Jette, C. A.; Abernathy, M. E.; Dam, K.-M. A.; Esswein, S. R.; Gristick, H. B.; Malyutin, A. G.; Sharaf, N. G.; Huey-Tubman, K. E.; Lee, Y. E.; Robbiani, D. F.; Nussenzweig, M. C.; West, A. P., Jr.; Bjorkman, P. J., SARS-CoV-2 neutralizing antibody structures inform therapeutic strategies. Nature 2020, 588, 682.

4. Sulea, T.; Baardsnes, J.; Stuible, M.; Rohani, N.; Tran, A.; Parat, M.; Donates, Y. C.; Duchesne, M.; Plante, P.; Kour, G.; Durocher, Y., Structure-based dual affinity optimization of a SARS-CoV-1/2 cross-reactive single-domain antibody. Plos One 2022, 17.

